# The *tidyomics* ecosystem: Enhancing omic data analyses

**DOI:** 10.1101/2023.09.10.557072

**Authors:** William J. Hutchison, Timothy J. Keyes, The tidyomics Consortium, Helena L. Crowell, Jacques Serizay, Charlotte Soneson, Eric S. Davis, Noriaki Sato, Lambda Moses, Boyd Tarlinton, Abdullah A. Nahid, Miha Kosmac, Quentin Clayssen, Victor Yuan, Wancen Mu, Ji-Eun Park, Izabela Mamede, Min Hyung Ryu, Pierre-Paul Axisa, Paulina Paiz, Chi-Lam Poon, Ming Tang, Raphael Gottardo, Martin Morgan, Stuart Lee, Michael Lawrence, Stephanie C. Hicks, Garry P. Nolan, Kara L. Davis, Anthony T. Papenfuss, Michael I. Love, Stefano Mangiola

## Abstract

The growth of omic data presents evolving challenges in data manipulation, analysis, and integration. Addressing these challenges, Bioconductor^1^ provides an extensive community-driven biological data analysis platform. Meanwhile, tidy R programming^2^ offers a revolutionary standard for data organisation and manipulation. Here, we present the *tidyomics* software ecosystem, bridging Bioconductor to the tidy R paradigm. This ecosystem aims to streamline omic analysis, ease learning, and encourage cross-disciplinary collaborations. We demonstrate the effectiveness of *tidyomics* by analysing 7.5 million peripheral blood mononuclear cells from the Human Cell Atlas^3^, spanning six data frameworks and ten analysis tools.

## Main

High-throughput technologies for genomics, epigenomics, transcriptomics, spatial analysis, and multi-omics have revolutionised biomedical research, presenting opportunities and challenges for data manipulation, exploration, analysis, integration, and interpretation^4^. To address these challenges, the scientific community has developed object-oriented frameworks for data organisation and specialised operations.

In response to the complexity of the software landscape, the Bioconductor Project^1^ has emerged as a premier R software repository and platform for omic data analysis. Bioconductor provides international standardisation and interoperability for data processing workflows and statistical analysis. With extensive annotation resources and standardised data formats that link metadata, Bioconductor promotes reproducibility and community-driven open-source development.

Recently, the tidy R paradigm and the tidyverse software ecosystem^2^ have transformed R-based data science by prioritising intuitive data representation and manipulation over complex data structures and syntax. This paradigm uses tables to represent data, with variables as columns and observations as rows. It simplifies data manipulation with operations connected in pipelines that use standardised and natural language vocabulary. The components of the tidyverse rank as the most frequently downloaded R packages^5^ and are widely taught in Data Science and Bioinformatics programs worldwide^6^.

Bioconductor has remained largely independent of the tidyverse ecosystem. Creating a bridge between these two ecosystems by providing a tidy interface to standard data formats^7–9^ and analysis^9^ would enable researchers to shift their focus from technical challenges to biological questions. Also, leveraging a standard in data science education would lower the barrier to entry for analysing diverse omic data.

Here, we present *tidyomics*, an interoperable software ecosystem that bridges Bioconductor and other omic analysis frameworks (e.g. Seurat^8^) with the tidyverse. This ecosystem is installable with a single homonymous meta-package, available in Bioconductor. *tidyomics* includes three new packages: tidySummarizedExperiment, tidySingleCellExperiment, and tidySpatialExperiment, and five publicly available R packages: plyranges^7^, nullranges^10^, tidyseurat^8^, tidybulk^9^, and tidytof^11^. Importantly, while *tidyomics* represents omic data in a tidy format **(Figure 1A)**, it leaves the original data containers and methods unaltered, ensuring compatibility with existing software, maintainability and long-term Bioconductor support.

**Figure 1:**
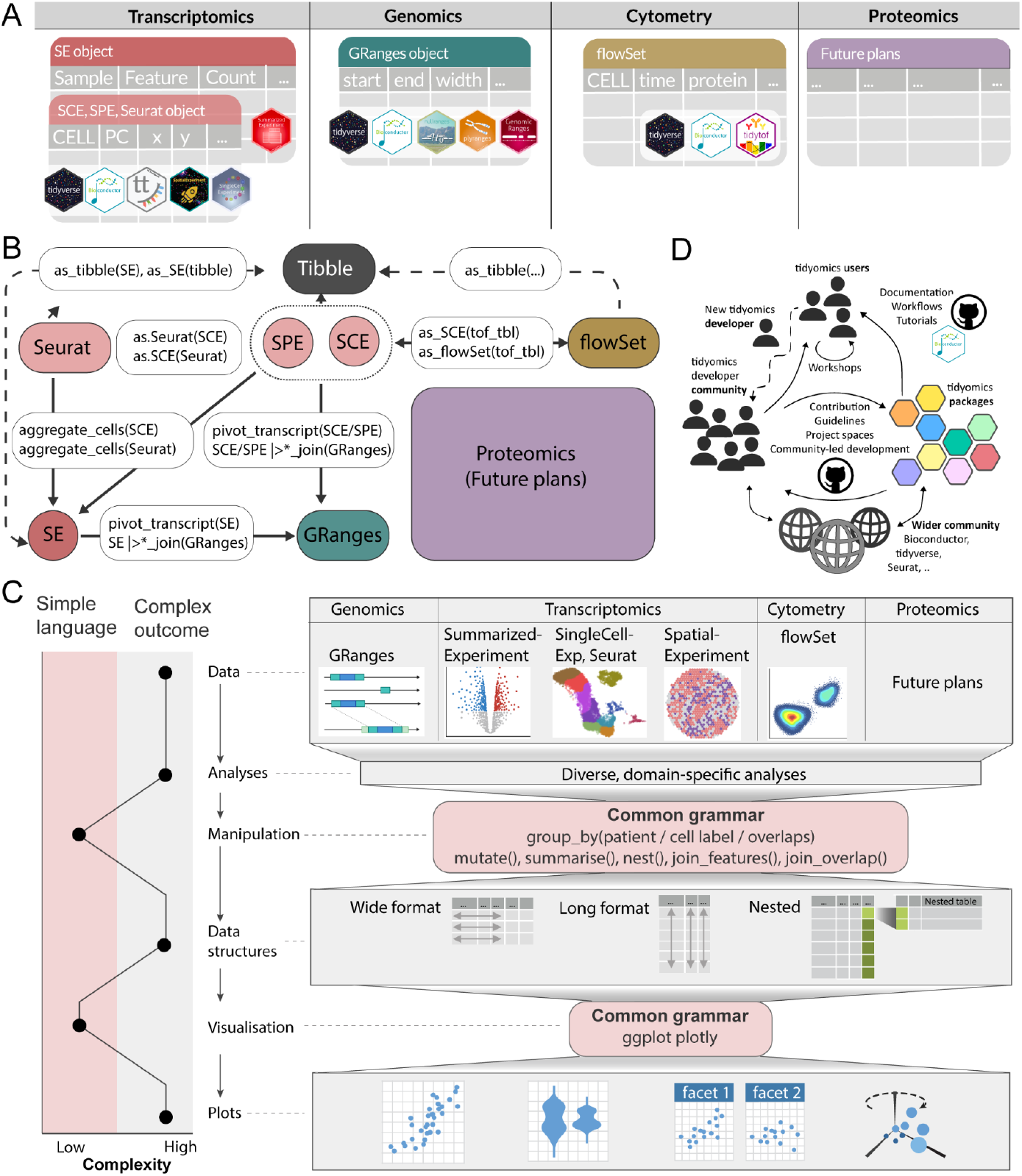
Overview of the *tidyomics* ecosystem. **A:** Diagrams of data interfaces show consistent data representation for the diverse data containers. The hexagonal icons represent the compatible R packages for each data container. **B:** The landscape of rich data objects in R/Bioconductor, with *tidyomics* verbs as paths connecting these objects. The data containers are represented by rounded rectangles and functions that connect them as white boxes. SPE = SpatialExperiment, SCE = SingleCellExperiment, SE = SummarizedExperiment. **C:** Contrast between the simplicity of the tidy syntax/grammar and the complex outcome and input data containers (left). Example workflows include data, biological analysis, data/results manipulation and summarisation, diverse data structures, visualisation and resulting plots (right). The pink areas include the infrastructure that shares grammar across omics. **D:** Engagement within the *tidyomics* community is multifaceted, centring around a suite of R packages tailored for streamlined data analysis. This ecosystem is enhanced by comprehensive documentation, including usage guidelines and development tutorials. The community thrives on interactive learning, offering workshops created and led by its members. Collaboration, project development and guidelines are centralised in our *tidyomics* GitHub organisation. GitHub and Bioconductor are the primary discussion forums. Additionally, Bioconductor is a prominent repository for software packages, reinforcing the community’s connection to broader bioinformatics networks such as Bioconductor, tidyverse, and Seurat.

Within Bioconductor, GenomicRanges^12^ organises genomic features as ranges (e.g. genes, exons, SNPs, CpGs) in rows, linked with variables (e.g. range width) as columns (like the BED^13^ format). plyranges^7^ extends dplyr verbs to GenomicRanges objects, facilitating ranges integration, overlap analysis, summarisation, and visualisation. plyranges interacts with nullranges for matching^14^ or bootstrapping^10^ ranges to perform overlap enrichment analyses.

In Bioconductor, SummarizedExperiment and SingleCellExperiment^15^ organise transcriptional abundance as a feature/gene-by-sample matrix linked with metadata. *tidyomics* generalises the concept of the *variable* by providing a tabular interface with observations (e.g. gene/sample pair) as rows and variables (e.g. abundance, metadata) as columns. This approach enables complex filtering, summarisation, analysis, and visualisation using the tidyverse. tidybulk^9^ offers a tidy and modular analysis pipeline for bulk and pseudobulk data.

Bioconductor’s flowCore^16^ package organises data from mass, flow and sequence-based cytometry in a cell-by-feature matrix and facilitates data manipulation. tidytof^11^ interfaces flowCore with the tidyverse, tidySingleCellExperiment, and tidySummarizedExperiment.

Bioconductor’s SpatialExperiment^17^ organises data from cell/pixel-based technologies^18^, such as 10X Genomics Xenium, CosMX, Mibi, and MERSCOPE. tidySpatialExperiment offers a tidy interface for data with spatial coordinates and provides specialised operations such as gating based on geometric and hand-drawn shapes.

*tidyomics* is a unified and interoperable software ecosystem for omic technologies that covers several omic analysis frameworks. Through conversion and join operations, a network of functionalities connects all data containers **(Figure 1B)**. This harmonised approach facilitates seamless container switching, decreasing the dependence on a specific framework created by domain-specific syntax and effectively increasing the umbrella of used tools **(Figure 1C)**.

To demonstrate *tidyomics’* utility and scalability, we tested sex transcriptomic differences of the peripheral immune system across 7.5 million blood cells. Our ecosystem seamlessly bridged six data and analysis frameworks **(Figure 2A)**, showcasing the benefit of consistently using tidy R grammar instead of mixing the syntaxes of base R, DuckDB, Seurat, SingleCell- and SummarizedExperiment, DGEList and GenomicRanges. After preprocessing, we tested 15,494 pseudobulk samples across 26 immune cell types **(Figure 2B)** with a multilevel differential expression model. We identified T CD4 naive cells, T effectors, and B memory cells as the most changing between sexes **(Figure 2C)**. Most sex-related transcriptional changes (excluding sex chromosomes) were cell-type specific rather than shared (FDR < 0.05) **(Figure 2D)**. We tested the proximity of genes with a significant effect for sex or its interaction with age in CD4 naive cells to GWAS SNPs for three immune-cell-related and sex-biased diseases: multiple sclerosis, rheumatoid arthritis, and systemic lupus erythematosus **(Figure 2E)**. We found nine genes overlapping or near SNPs associated with these diseases (*IL2RA, CD40*, and *KCP* associated with two or more). A large proportion of sex-related genes, 41%, define divergent immune-ageing trajectories **(Figure 2F)**.

**Figure 2:**
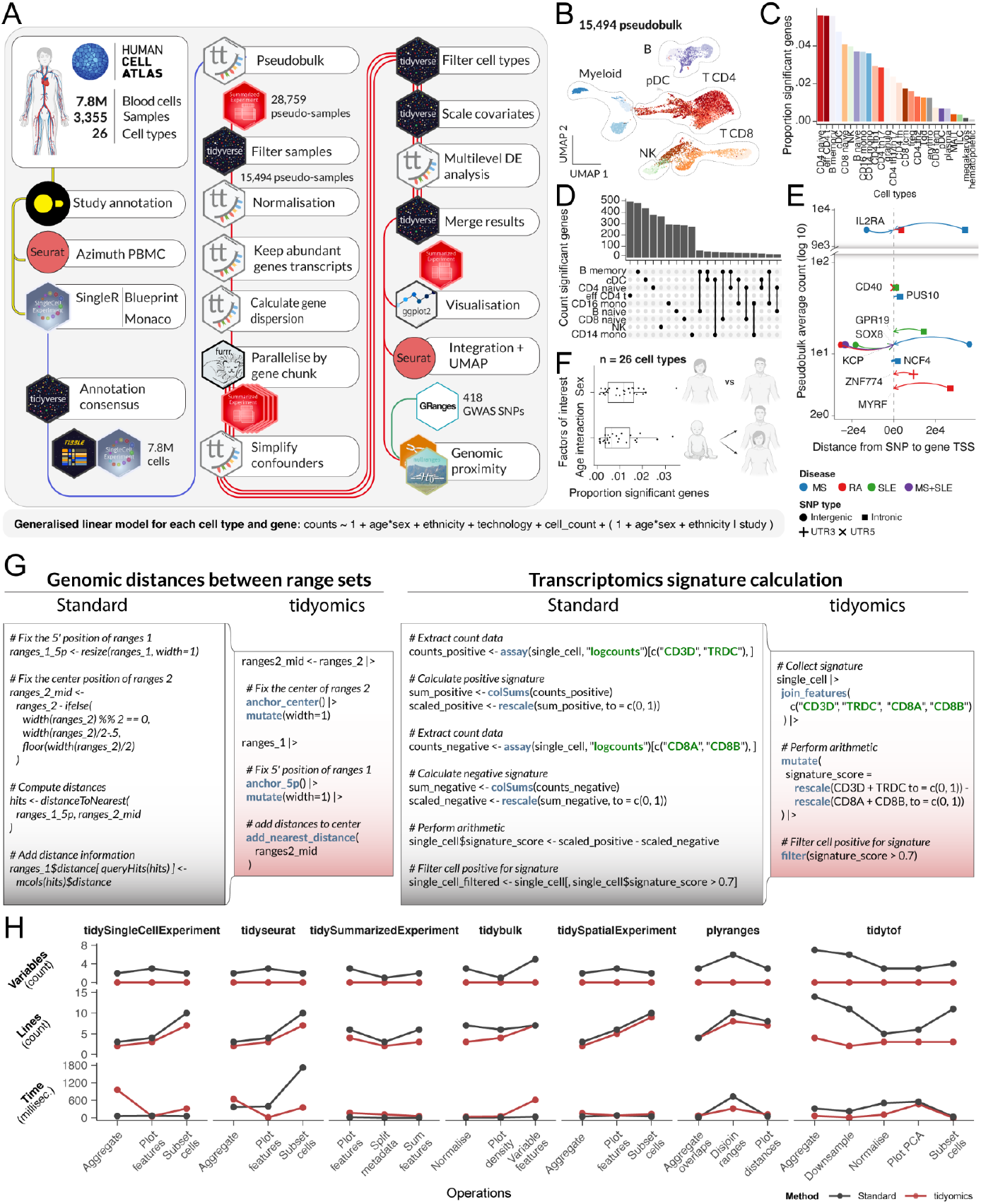
Performance of the *tidyomics* ecosystem. **A:** *tidyomics* powers large-scale cross-framework analyses. We compared peripheral blood mononuclear cells between sexes at the pseudobulk level. The logos represent data and analysis frameworks. The connecting lines represent pipelines, coloured by the object type. Parallel lines represent parallel workflows. **B:** Pseudosample UMAP, coloured by cell type. **C:** Rank of cell types from the most to the least changed across sexes, coloured as per the B panel. **D:** Significant gene overlap across the top nine cell types for sex effect or its interaction with age. **E:** Overlap of sex-related genes in CD4 naive cells with GWAS SNPs for multiple sclerosis, rheumatoid arthritis, and systemic lupus erythematosus. **F:** Fraction of sex-related genes significant as a main effect or interaction with age. The box plot centre line represents the median value, and the lower and upper hinges represent the first and third quartiles. The lower whisker extends from the lower hinge to 1.5 times the interquartile range or the lowest value. The upper whisker extends from the upper hinge to 1.5 times the inter-quartile range or the highest value. **G:** Comparison of code readability between standard and tidyverse programming. Two tasks showcased are visualising a histogram of genomic distances (left) and calculating a multi-gene signature from single-cell data (right). **H:** The benchmark of variables, lines of code, and time efficiency of our ecosystem compared to standard (non-tidy) coding. The operations include common manipulations and analysis for each package (Methods).

This analysis shows that *tidyomics* allows code repurposing across diverse data sources. For example, complex manipulation and visualisation of genome and transcriptome data can be performed using modular, consistent grammar assembled into a compact and legible pipeline **(Figure 2G)**. Legibility and coding simplicity are promoted by fewer intermediate variables and lines of code compared with standard counterparts while incurring no major execution-time overhead **(Figure 2H)**. Tidy R favours functional programming (e.g., vectorisation rather than for-loops), which helps avoid bugs caused by variable updating, especially in interactive programming.

The Bioconductor coding standards and contribution guidelines adopted by our dedicated developer community set a solid ground for the long-term maintainability of the ecosystem **(Figure 1D)**. This ecosystem will grow, including R packages with compatible goals and standards from Bioconductor and CRAN. While *tidyomics* is currently focused on simplification and harmonisation, novel analysis and manipulation tools are among its goals.

The tidyverse and Bioconductor ecosystems are transforming R-based data science and biological data analysis. *tidyomics* bridges the gap between these ecosystems, enabling analysts to leverage the power of tidy data principles in omic analyses. This integration fosters cross-disciplinary collaborations, reduces barriers to entry for new users, and enhances code readability, reproducibility, and transparency. The tidy standard applied to biological software creates an extensible development ecosystem where independent researchers can interface new software. Ultimately, the *tidyomics* ecosystem, consisting of new and publicly available R packages, has the potential to significantly accelerate scientific discovery.

## Acknowledgements

We acknowledge Bioconductor and tidyverse communities, whose software and coding paradigms this work is based on and would not be possible without. We also thank the *tidyomics* community for their feedback and contribution. We thank Vincent Carey for his support and feedback on the project. Also, we thank Matt Richie for his continuous support and feedback. Human illustrations were created with BioRender.com.

S.M. was supported by the Victorian Cancer Agency Early Career Research Fellowship (ECRF21036). M.I.L. was supported by the Chan Zuckerberg Initiative (EOSS3-0000000057). A.T.P. was supported by the National Health and Medical Research Council (NHMRC) Senior Research Fellowship (1116955) and Investigator Grant (2026643). A.T.P., S.M., and W.H. were supported by the Lorenzo and Pamela Galli Medical Research Trust and the Galli Next Generation Discoveries Initiative. K.L.D. is the Anne T. and Robert M. Bass Endowed Faculty Scholar in Pediatric Cancer and Blood Diseases of the Stanford Maternal Child Health Research Institute and the Harriet and Mary Zelencik Endowed Faculty in Children’s Cancer and Blood Diseases. P.P.A. was supported by the Cancéropole GSO and Intergroupe Français du Myélome. R.G. was funded by a project grant from the Swiss National Foundation. M.M. was supported by the NHGRI and NCI of the National Institutes of Health under award numbers U41HG004059 and U24CA180996. This work was supported by an ASPIRE award from the Mark Foundation for Cancer Research and the B+ Foundation. The research benefitted from support from the Victorian State Government Operational Infrastructure Support and Australian Government NHMRC Independent Research Institute Infrastructure Support. The funders had no role in study design, data collection and analysis, or decision to publish or prepare the manuscript.

## Author information

### Authors Contributions

S.M. proposed the study, S.M. and M.I.L. designed the study. W.J.H. and S.M. developed the novel tidy adapters for transcriptomics, W.J.H., T.J.K., S.M., and M.I.L. performed the analyses. W.J.H., T.J.K., H.L.C., J.S., C.S., E.S.D., N.S., L.M., B.T., A.A.N., M.K., Q.C., V.Y., W.M., J.E.P., I.M., M.H.R., P.P.A., P.P., C.L.P., M.T., R.G., M.M., S.L., M.L., S.C.H., G.P.N., K.L.D., A.T.P., M.I.L., and S.M. contributed to the ecosystem’s development and ongoing improvement. S.M., M.I.L., A.T.P., K.L.D., S.C.H., M.L., M.M., and R.G. acted as the supervisory team. S.M., M.I.L., and A.T.P. contributed equally and jointly led the study. W.J.H. and T.J.K. contributed equally. All authors contributed to the manuscript’s writing.

## The tidyomics Consortium

William J. Hutchison ^1, 2^, Timothy J. Keyes ^3, 4^, Helena L. Crowell ^5, 6^, Jacques Serizay ^7^, Charlotte Soneson ^8, 9^, Eric S. Davis ^10^, Noriaki Sato ^11^, Lambda Moses ^12^, Boyd Tarlinton ^13^, Abdullah A. Nahid ^14^, Miha Kosmac ^15^, Quentin Clayssen ^16^, Victor Yuan ^17^, Wancen Mu ^18^, Ji-Eun Park ^18^, Izabela Mamede ^19^, Min Hyung Ryu ^20, 21^, Pierre-Paul Axisa ^22^, Paulina Paiz ^3^, Chi-Lam Poon ^23^, Ming Tang ^24^, Raphael Gottardo ^25, 26, 9^, Martin Morgan ^27^, Stuart Lee ^28^, Michael Lawrence ^28^, Stephanie C. Hicks ^29, 30, 31^, Garry P. Nolan ^32^, Kara L. Davis ^4^, Anthony T. Papenfuss ^1, 2^, Michael I. Love ^33, 18^, Stefano Mangiola ^1, 2, 34^

## Methods

### tidySummarizedExperiment

tidySummarizedExperiment introduces an innovative and versatile approach for representing and managing bulk data, offering an alternative to and complementing the conventional methodologies commonly employed in SummarizedExperiment. This novel approach incorporates adaptors tailored to widespread data manipulation and visualisation packages, including dplyr, tidyr, ggplot, and plotly. Crucially, the core structure of the SummarizedExperiment object remains unaltered, ensuring seamless compatibility with existing analytical workflows.

Data is represented as a long table, wherein observations are defined by the abundance of a feature/sample pair, while variables encompass feature and sample-related metadata. The fundamental columns comprising this representation consist of the feature and sample columns. Importantly, when any of these essential columns are absent from the output of a given operation (e.g., select) or when a summarised version of these columns is generated, an independent table is returned. This supplementary table adheres to the structure of the SummarizedExperiment tidy representation, facilitating separate analysis or visualisation as needed. Furthermore, when the returned subset of observations does not represent a valid SummarizedExperiment (e.g. it does not correspond to a rectangular slice of the feature/sample matrix), a tibble is returned for independent analyses.

With the tidySummarizedExperiment approach, newly created or joined columns, such as those obtained from a metadata table, are automatically incorporated into the colData, rowData, or assays based on their alignment with feature or sample identifiers. This versatile mechanism extends the “variable” concept, enabling a manipulation, displaying, or visualisation of sample, feature, and abundance information with consistent grammar. Notably, the columns for the sample and feature identifiers and genomic ranges are designated as read-only to preserve data integrity.

### tidySingleCellExperiment

tidySingleCellExperiment presents a novel approach for representing and manipulating single-cell data, providing an alternative to and complementing the conventional methods commonly used in SingleCellExperiment. The main goal and property of the API are consistent with tidySummarizedExperiment.

However, as the central analysis unit of single-cell data is cells, rather than genes for bulk data, tidySingleCellExperiment favours the cell-wise (metadata and reduced dimensions, e.g., principal components and UMAP dimensions) and sample-wise information rather than gene-wise and sample-wise information for tidySummarizedExperiment. This design choice keeps the single-cell data representation highly interpretable and practically useful at the cost of a partial lack of consistency to the bulk data. The emphasis on cell-wise information over transcript-wise information is driven by the priority to facilitate ease of use, data summarisation, information integration, and data visualisation in the context of cell analysis and by explicit feature-wise operation not being as common as cell-wise operation. By focusing on cell-wise information, the abstraction avoids unnecessary complexity from including feature-level information (e.g., genes, proteins, genomic regions) by default, especially when performing cell-wise information subsetting.

To access transcript-level information, users can utilise the ‘join_features’ function. This function enriches the metadata by incorporating transcript identifiers, transcript abundance, and transcript-wise annotations, including gene length, genomic coordinates, and functional annotations, as additional columns of the cell metadata.

The tibble abstraction employed in tidySingleCellExperiment consists of two column types: editable columns, which allow user interaction and modification, and view-only columns, which encompass data-derived variables, such as reduced dimensions. Integrating all cell-wise information, including reduced dimensions, within a single tibble representation enables seamless data visualisation, filtering, and manipulation. Importantly, this design ensures compatibility with the tidyverse ecosystem, enabling data manipulation and plotting using routines from dplyr, tidyr, ggplot2, and plotly. Furthermore, adopting this abstraction allows users to operate on the data as if it were a standard tibble while preserving compatibility with any other algorithms or tools that utilise the SingleCellExperiment framework. This approach ensures full backward compatibility and facilitates seamless integration into existing workflows.

tidySingleCellExperiment shares the same grammar and data representation as tidyseurat, allowing users to use tidy code with SingleCellExperiment and Seurat data containers.

### tidySpatialExperiment

tidySpatialExperiment provides a tidy R abstraction (tibble) of SpatialExperiment objects. Similarly to tidySingleCellExperiment, it provides cell-wise information, including cell metadata, spatial coordinates and metadata, and reduced dimensions. All information can be processed with tools provided by dplyr, tidyr, ggplot2 and plotly. tidySpatialExperiment provides the ‘join_features’ function to append the specified features to the cell-wise information and, consequently, the tibble abstraction. The ‘aggregate_cells’ function is provided to combine cells by shared variables and aggregate feature counts.

### Benchmarking

We benchmarked standard and tidyomics workflows for common data analysis tasks. Briefly, the aggregate-cells-by-sample operation aggregates the feature counts of cells within each sample. The plot features per cell operation plots the distribution of summed feature counts for each cell within each sample. The subset cells by feature operation subsets cells by feature signature threshold. The subset-features-by-mean-count operation subsets features by mean count threshold. The plot-features-per-sample operation plots the distribution of feature counts for each sample. The normalise-features operation normalises feature counts across samples. The plot-normalised-feature-density operation plots the density of normalised features for each sample. The identify-variable-features operation identifies the most variable features for each cell type. The aggregate-overlaps operation identifies and aggregates overlapping regions. The plot-feature-set-distances operation calculates and plots the distance between feature sets. The group-disjoin-ranges operation finds disjoint regions within groups of features and subsets overlaps. The downsample-cells operation randomly subsets cells from each sample. The plot-PCA operation calculates and plots principal components. The benchmarking operations were run using R v4.3.1, plyranges v1.22.0, tidySingleCellExperiment v1.13.3, tidySpatialExperiment v0.99.13, tidySummarizedExperiment v1.12.0, tidybulk v1.15.4, tidytof v0.0.0, and tidyseurat v0.8.0. Each benchmarking operation was executed 50 times using the microbenchmark package, and the mean time elapsed was recorded. The variable count was calculated as the number of times a new variable was created to store data. The line count was calculated as the number of lines required for each operation while following indentation best practice.

### Transcriptional analyses

We collected all human peripheral blood mononuclear cells from the Human Cell Atlas^1^ initiative using the CELLxGENE database. We downloaded the metadata and gene-transcript abundance through the R package CuratedAtlasQuery (10.18129/B9.bioc.CuratedAtlasQueryR). We consistently represented age as days (named ‘age_days’ in our database). Where a categorical value was provided, we converted it into equivalent days using publicly available references. For example, the “adolescent stage” was converted to 15 years old (= 5475 days).

Immune cells were labelled using Seurat Azimuth mapping to the PBMC reference^2^ (using tidyseurat^3^) and SingleR^4^ with the Blueprint^5^ and Monaco^6^ references^4^. To identify a consensus, we compared and contrasted the high-resolution labels ‘predicted.celltype.l2’ for Seurat Azimuth and ‘label.fine’ Blueprint and Monaco references. When possible, the reference-specific cell-type labels were standardised under a common ontology (Table SX). Where the resolution of transcriptionally similar cell types was uncertain with the given tools, cell types were labelled with a coarser resolution. For example, innate lymphoid and natural killer cells were grouped under “innate lymphoid”. The cell type curation was performed to obtain a high confidence, meaningful representation of the immune system’s heterogeneity, allowing data-rich cell types whose tissue composition can be modelled probabilistically rather than aiming for the finest resolution possible. The original annotation provided by the studies was integrated with the three new annotation sources to identify a total or partial consensus. Cell types without complete or partial consensus were filtered out.

We selected primary (no re-analysed data or collections) physiological samples (i.e. no disease). We also excluded samples with less than 30 cells. We excluded erythrocytes and platelets from the analyses as they were not of interest. To increase the sample size per demographic group, we merged Asian descendants (labelled in CELLxGENE as Asian and Chinese). We excluded samples whose ages were unknown.

tidySingleCellExperiment aggregated cells across samples and cell types in pseudobulk transcript counts. Pseudobulk samples with less than 5000 genes or 10 cells were filtered out using tidySummarizedExperiment. Quantile normalisation in Limma^7^ was used through tidybulk^8^, as across the 28,241 pseudo samples, the data distribution was heterogeneous and non-controllable. Lowly abundant gene transcripts were filtered out using edgeR^9^ through tidybulk, using sex and ethnicity as factors of interest, minimum counts of 500, and minimum proportion of 0.9.

Gene-transcript abundance for each cell type was modelled using the following formulation, with age as a centred and scaled continuous variable (mean age of ∼47 years).

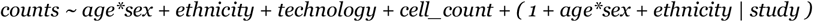

Given the complexity of the model, tidybulk was also used to identify data subsets that included complete covariate confounders. Tidybulk was used to fit the multilevel model through glmmSeq^10^ and test hypotheses (FDR < 0.05). Sex-related transcriptional changes (Figure 2C-F) were defined as genes significant for the main effect of sex or its interaction with age, excluding genes on sex chromosomes.

Seurat^11^ and tidyseurat^3^ were used to remove the study effect across cell types from the pseudobulk data and calculate the UMAP dimensions. Ggplot2 was used for visualisation^12^.

To overlap genes with significant effect for sex or its interaction with age in CD4 naive cells with GWAS lead SNPs, we used the tidySummarizedExperiment and plyranges packages to harmonise summary statistics from pseudobulk DE analysis and three GWAS for multiple sclerosis^13^, rheumatoid arthritis, and systemic lupus erythematosus^14^. As the GWAS data was provided in all cases for the hg19 genome build, the gene locations were loaded from the Bioconductor package TxDb.Hsapiens.UCSC.hg19.knownGene. The overlap analysis used genes with either significant main effect for sex or sex * age, as estimated from the pseudobulk multilevel model, with FDR < 0.05. Overlap was calculated as GWAS SNPs within 50 kb from the gene body, and distance was calculated from the GWAS lead SNP to the gene’s TSS.

## Data availability

Human Cell Atlas peripheral blood mononuclear single-cell data was downloaded from the CELLxGENE database (https://cellxgene.cziscience.com/). Metadata and gene-transcript abundance for these datasets were downloaded via the CuratedAtlasQuery R package (available in Bioconductor). CELLxGENE sample accession codes are available in the supplementary material. Source data for Figure 2H is available in the supplementary material.

## Code availability

The tidyomics homepage is https://github.com/tidyomics, which provides links to the constituent packages. The *tidyomics* meta-package is available at Bioconductor bioconductor.org/packages/tidyomics/. The tidySummarizedExperiment package is available at Bioconductor bioconductor.org/packages/tidySummarizedExperiment. The tidySingleCellExperiment package is available at Bioconductor bioconductor.org/packages/tidySingleCellExperiment. The tidySpatialExperiment package is available at Bioconductor bioconductor.org/packages/tidySpatialExperiment/. The code used to benchmark workflow efficiency and analyse peripheral blood mononuclear cells from the Human Cell Atlas is available at github.com/tidyomics/tidyomics_paper.

